# Traveling waves in the prefrontal cortex during working memory

**DOI:** 10.1101/2021.04.08.438959

**Authors:** Sayak Bhattacharya, Scott L. Brincat, Mikael Lundqvist, Earl K. Miller

## Abstract

Neural oscillations are evident across cortex but their spatial structure is not well-explored. Are oscillations stationary or do they form “traveling waves”, i.e., spatially organized patterns whose peaks and troughs move sequentially across cortex? Here, we show that oscillations in the prefrontal cortex (PFC) organized as traveling waves in the theta (4-8Hz), alpha (8-12Hz) and beta (12-30Hz) bands. Some traveling waves were planar but most rotated. The waves were modulated during performance of a working memory task. During baseline conditions, waves flowed bidirectionally along a specific axis of orientation. Waves in different frequency bands could travel in different directions. During task performance, there was an increase in waves in one direction over the other, especially in the beta band.

**Author Summary:** We found that oscillations in the prefrontal cortex form “traveling waves”. Traveling waves are spatially extended patterns in which aligned peaks of activity move sequentially across the cortical surface. Some traveling waves were planar but most rotated. The prefrontal cortex is important for working memory. The traveling waves changed when monkeys performed a working memory task. There was an increase in waves in one direction over the other, especially in the beta band. Traveling waves can serve specific functions. For example, they help maintain network status and help control timing relationships between spikes. Given their functional advantages, a greater understanding of traveling waves should lead to a greater understanding of cortical function.

## Introduction

Oscillatory dynamics have been linked to a wide range of cortical functions. For example, higher frequency (gamma, >40 Hz) power (and spiking) increases during sensory inputs (and their maintenance) and during motor outputs (1–5). Gamma power is anti-correlated with lower frequencies (alpha/beta, 8-30 Hz), whose power is often higher during conditions requiring top-down control (e.g., when attention is directed away, or an action is inhibited) (6–8). Such observations have led to a theoretical framework in which oscillatory dynamics regulate neural communication (9–11).

Thus far, most studies of neural oscillations have focused on what we can call “standing wave” properties (e.g., power of and coherence between oscillations at different cortical sites), ignoring any organization of where and when the peaks and troughs of activity appear. However, there is mounting evidence for such organization. Oscillations can take the form of “traveling waves”: Spatially extended patterns in which aligned peaks of activity move sequentially across the cortical surface (12,13). This apparent movement of the amplitude peak is facilitated by the existence of a phase gradient along a particular direction, along which the movement occurs. Traveling waves have most often been reported in the lower-frequency bands (<30 Hz). Examples include beta-band (15-30 Hz) traveling waves in motor and visual cortices (14,15) and theta band (3-5 Hz) traveling waves in the hippocampus (16,17).

Traveling waves are of interest because they have a variety of useful properties for cognition, development, and behavior. They can create timing relationships that foster spike-timing-dependent plasticity and memory encoding (14,18). They add information about recent history of activation of local networks (19). They are thought to help “wire” the retina (20) and cortical microcircuits during development (21). Their functional relevance in the adult brain is suggested by observations that traveling wave characteristics can be task-dependent and that they impact behavior. For example, EEG recordings have shown that alpha band waves reverse their resting state direction during sensory inputs (22). Behavioral detection of weak visual targets improves when there are well organized low-frequency (5-40Hz) traveling waves in visual cortex vs when there is a weaker, “scattered” organization (23).

Oscillatory activity in the prefrontal cortex has been linked with cognitive functions like working memory and attention but there has been little examination of whether they form spatio-temporal structures like traveling waves. Most studies have averaged oscillations across spatially distributed electrodes. This increases the “signal” of the oscillations but prohibits analysis of any spatial organization. Thus, we examined their spatial organization from microarray recordings in the PFC of monkeys performing a working memory task. This revealed that low-frequency (beta and lower) traveling waves are common in the PFC. We characterized their speed, direction and their patterns. They often rotated and changed direction during performance of a working memory task.

## Results

Two animals performed a delayed match-to-sample task (Figure 1A). They maintained central fixation while a sample object (one of eight used, novel for each session) was briefly shown. After a two-second blank memory delay (with maintained fixation), two different objects (test screen, randomly chosen from the eight) were simultaneously presented at two extrafoveal locations. Then the animals were rewarded for holding fixation on the object that matched the sample. Two 8×8 multi-electrode “Utah” arrays, one in each hemisphere, were used to record local field potentials (LFPs) from the dorsal pre-frontal cortex (dlPFC). All data is from correctly performed trials. We analyzed data from 14 experimental sessions, five from one animal (Animal/Subject 1) and nine from the other (Animal/Subject 2).

**Figure 1:**
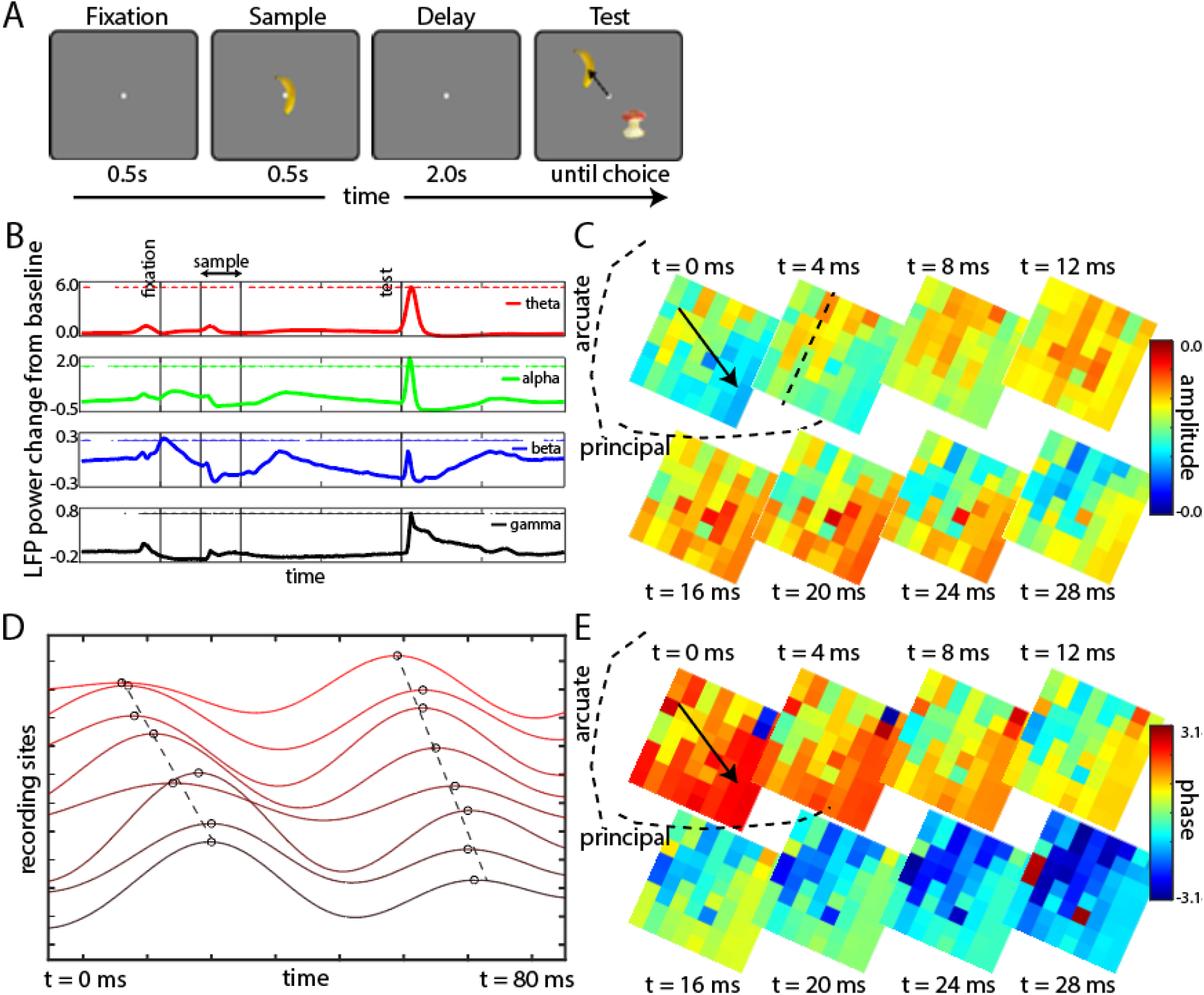
(A) Delayed-match-to-sample (DMTS) working memory task. The subject fixated at the center for 0.5 s (leftmost panel) before an object (one of eight) were presented at the center for another 0.5 s. There was a 2 s blank memory delay after which a test screen was presented at extrafoveal locations. The subject had to saccade towards the remembered object and hold fixation there (arrow, rightmost panel). (B) Filtered LFP power trends for all electrodes (four arrays), trial-averaged and shown across time with the lines denoting the major trial epochs. Four frequency ranges were chosen – theta (4-8 Hz), alpha (8-12 Hz), beta (12-30 Hz) and gamma (40-120 Hz), with the dots above the curve denoting if the LFP power at that instant is significantly (p<0.01) different from baseline (0.5 before fixation started). (C) Theta LFP organization across the right hemisphere array of Subject 1 of a particular trial. Each tile denotes a recording site on the array (8×8 total) with the color denoting the LFP amplitude at that site. The array position with respect to anatomical brain landmarks is overlaid. Each panel denotes the organization at an instant in time. The black arrow in the second panel indicates the direction of movement of the high amplitude LFPs with time. (D) Voltage traces from the 8 adjacent electrodes (color graded by position) in a row of the array in (C – dashed line). The circles mark the peak positions of the oscillation cycles with the dashed line indicating how the peaks shift gradually in time and space. (E) Phase maps from the corresponding panels in (C) with the color on each tile denoting the phase of the LFP oscillation cycle on that recording site.

### LFP oscillations formed traveling waves

Task-related modulations of LFP power were similar to that seen in previous studies. Figure 1B shows average changes in LFP power from baseline (a 500ms period just before fixation) across all electrodes, all sessions, and both animals as a function of time during the trial. Alpha (8-12 Hz) and beta (12-30 Hz) power decreased during sample presentation and increased during the delay relative to baseline. By contrast, theta (4-8 Hz) and gamma (40-120 Hz) power showed opposite trends. They increased during sample presentation and test screen presentation. Gamma power also ramped up near the end of the delay.

These increases in oscillatory power were not equal or simultaneous across the electrode array. Rather, there was a spatio-temporal structure that suggested traveling waves of activity. An example of a beta-band (12-30 Hz) wave during test screen presentation of one trial is shown in Fig. 1C (Supplementary Materials - Movie 1). The tiles represent the electrodes in the array. The array is oriented relative to its position with respect to anatomical brain landmarks in the dlPFC. The colors show the beta-band signal amplitude at each electrode at different snapshots in time. Note that, at first (t=0), beta is higher near the top of the array. Then, over time, the higher beta amplitude shifts systematically to the bottom of the array. To illustrate this in another way, Figure 1D shows the filtered LFP traces across one row of the array (dashed line in Fig. 1C; the electrodes are plotted on the Y-axis from left to right ascending). The oscillation peaks are marked. A sequential shift of the LFP peak activation across the row of electrodes can be seen. This shifting peak of activity in space with time is characteristic of traveling waves.

As a traveling wave moves across cortex, it should produce a phase gradient across adjacent recording sites. This can be seen in Figure 1E. It replots the data from Figure 1C as a phase map. The instantaneous beta-band phase at each electrode is color-coded for each time slice. Zero phase is the peak of each oscillation. Positive values indicate LFPs that were on their way to peak amplitude. Negative values indicate LFPs that have passed the amplitude peak. Note the phase gradient across the array and how it changes over time. At first, positive phase values tend to be higher on the bottom of the array. Then, as the wave travels along the arrow, the phase values on the top become negative (indicating that they are past the peak of the wave), phase values in the bottom shift toward zero over time (indicating the wave approaching peak) followed by a shift to negative values. But, at each point in time, the phase gradient consistently points in the direction of wave propagation. This can also be seen in Figure 1D. At the timepoints when the topmost electrode is at peak phase, the rest of the electrodes form a gradient down the rising phase of the sinusoidal wave. Thus, the instantaneous phase gradient provides a signature of a traveling wave and its direction at each point in time during a wave traversal.

A traveling wave was counted when there was a strong phase gradient in a particular direction. We counted the number of traveling waves across all sessions and both arrays in both animals by computing the trial-by-trial correlation between the observed phase map at each time point and a Euclidean distance map (with increasing distance in the direction being checked, see Methods and Supplemental Fig. 1A). Quadrant-based distance maps were used to calculate waves across twelve directions (all pairwise combinations of four quadrants × two directions, see Methods). When the correlation between the distance and phase map exceeded a certain threshold, a wave in that direction was counted. This threshold was set such that detected waves were required to have correlations greater than 99% of those obtained by random shuffling of the electrode locations in the array (see Methods). We further used simulations to validate our wave classification algorithm (Methods). We applied this classification across time in each trial using a shifting 20ms time-interval (short enough to capture anything less than 50Hz), averaging across trials and across all four arrays in the two animals.

Waves were apparent throughout the trial, albeit at different times for different frequency bands. Figure 2A shows the wave count (summed across all directions) for the theta, alpha, and beta frequency bands. For all bands, there were consistent increases in wave counts, relative to baseline (a 500ms window before fixation), during fixation, sample and test screen presentations (Fig. 2A). They elicited bursts of sharp, consecutive increases and decreases in wave prevalence over time (i.e., significantly above and below baseline values corresponding to a time-locked traveling wave followed by brief pause).

**Figure 2:**
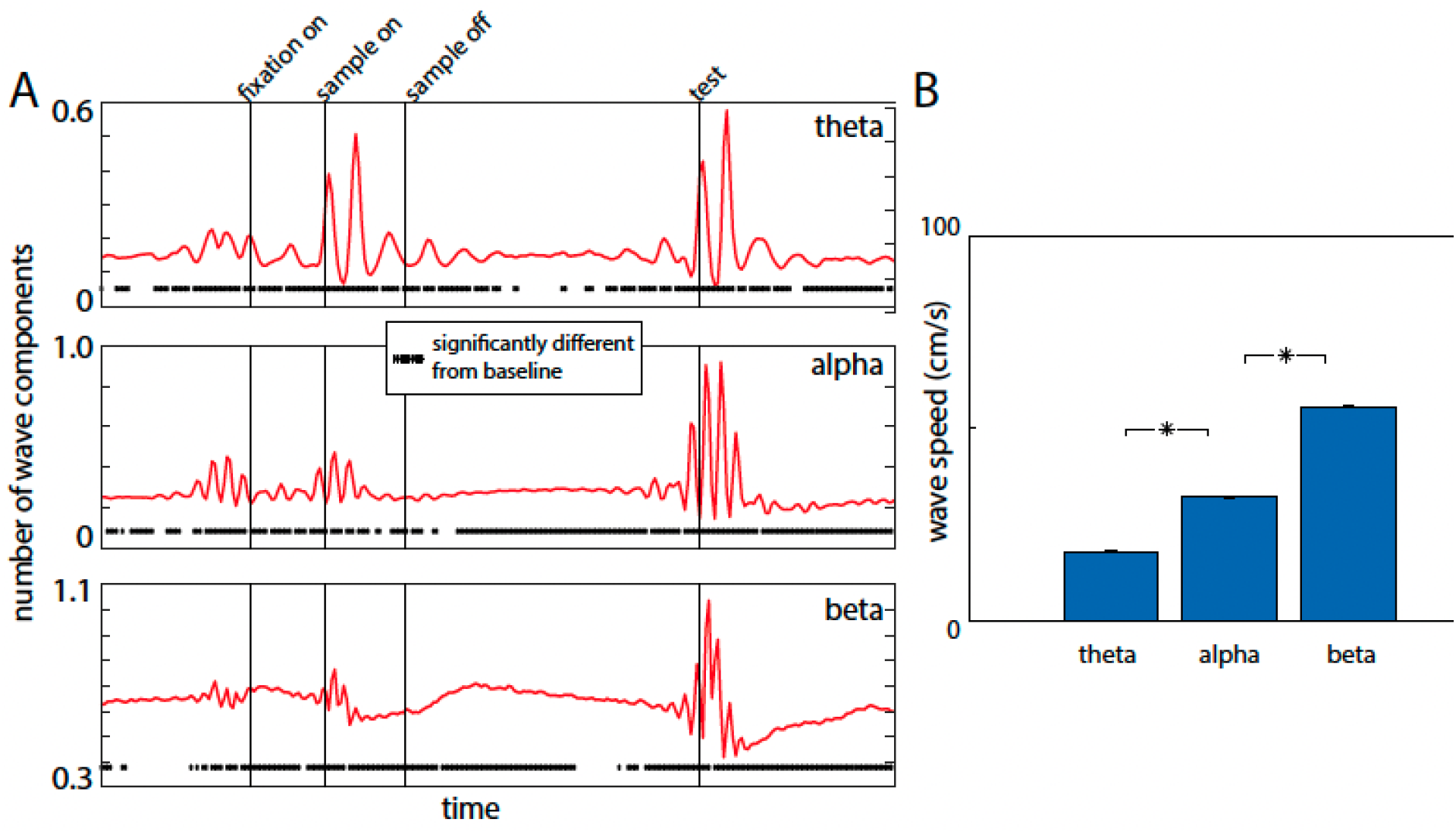
(A) Number of wave components observed in the three frequency ranges during the trial. The numbers are trial-averaged across all four arrays. The dots underneath denote if the numbers at that instant are significantly different from baseline (0.5s before fixation). (B) Wave speeds observed across all frequency ranges. Error bars represent SEM. Star indicates statistically significant difference (p<0.01).

The wave trends over time (Fig 2A) mirrored the power modulations shown in Fig 1B. For the theta and alpha bands, there were significantly higher-than-baseline wave counts in all epochs. Counts peaked during sample presentation (for theta) and test screen presentation (for theta and alpha). Beta waves also increased relative to baseline in all epochs. However, in contrast to alpha and theta bands, beta traveling waves tended to be higher during the memory delay and decreased during sample and test screen presentation. Wave speed ranged from 20-60 cm/sec and increased with increases in frequency band (Fig. 2B) as expected (15).

### Rotating waves

We found that the waves often had a rotational component. To determine this, we examined the circular-circular correlation coefficient (ρ_c_) between observed phase maps and rotation maps across different points of the array (see Methods). This represented the circular correlation between a rotation map centered around a point on the array and the array phase map for that slice of time. The correlation provided an estimate of the wave direction vector. For example, in Fig 3A, the red arrows showed positive coefficients for the central correlation coefficient (see Methods, Supplemental Fig. 1B).

**Figure 3:**
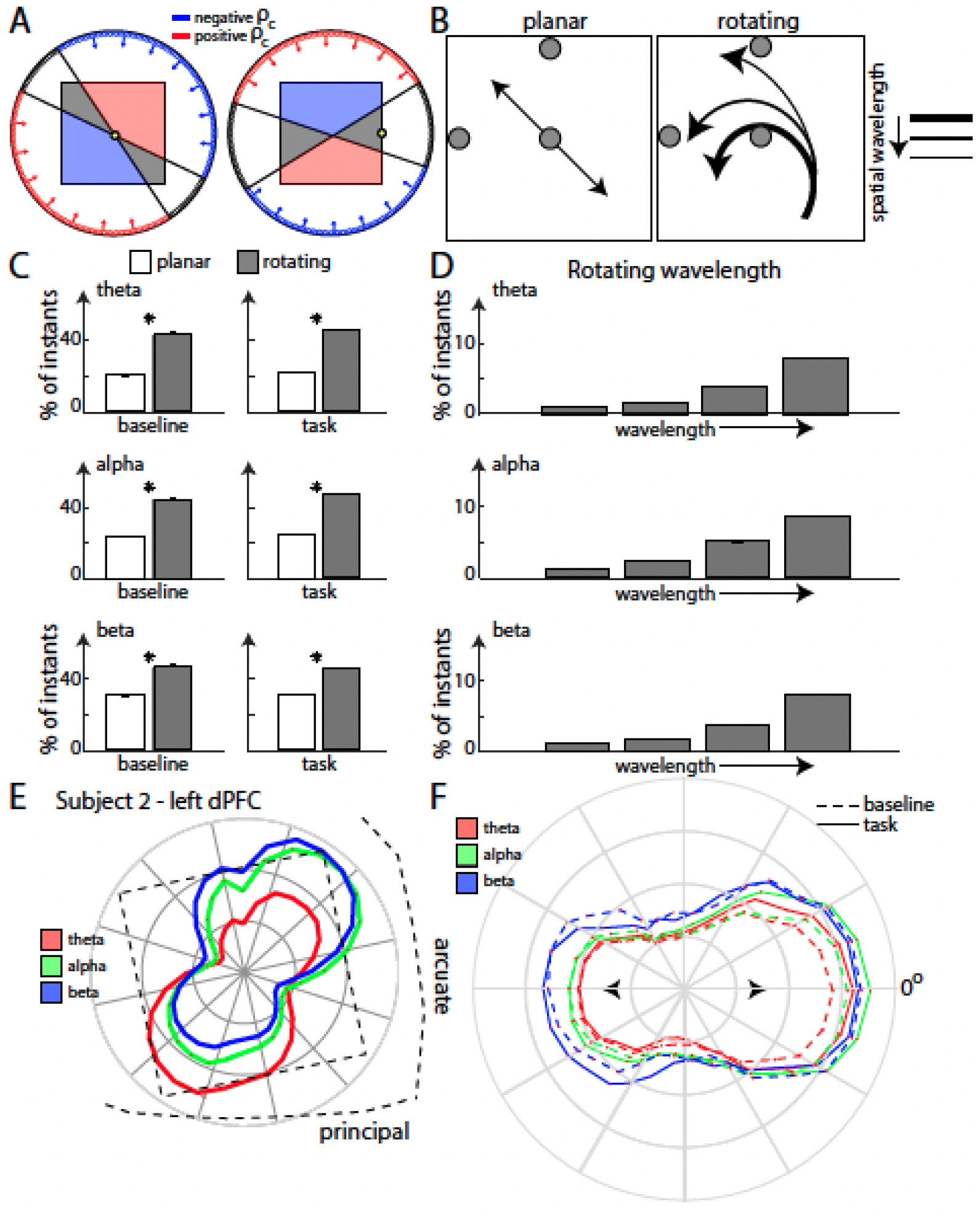
(A) Schematic illustrating the function of the circular-circular correlation coefficient (rho) around a particular point (yellow circle). Two different positions of the point are shown. A positive rho is calculated for the waves toward the red part of the array (red vectors), while blue gives a negative rho. The grey zone denotes wave directions that gave a correlation coefficient value within the “chance zone”. The zones differ with choice of point (right vs left panel) (B) Two broad wave types – planar (left) and rotating (right) waves were classified. Finer distinction possibilities are also shown – different directions (planar) or different spatial wavelengths (rotating). A combination of three coefficient values were used to distinguish between these wave types. The grey circles denote the points on the array around which the coefficient was calculated. (C) Plots showing the number of time instants in which rotating/planar waves were observed across all arrays and trials in the three frequency ranges. Star indicates statistically significant difference (p<0.01). (D) Plots showing the different wavelengths observed for the rotating waves classified in (C). (E) Wave directions (planar and rotating combined) observed across all trials in Subject 2 (left hemisphere) overlaid on the array position with respect to brain landmarks. The three colors denote the three frequency ranges. (F) Wave directions similar to (E), but for all four arrays combined, with each array’s most preferred direction aligned along the 0-degree axis.

A range of ρ_c_ captured a spectrum of wave types, from planar to rotating waves, of different directions and spatial wavelengths (Fig. 3B, see Methods). A high ρ_c_ value indicated a wave moving in a particular direction – but did not distinguish between the type of wave (planar vs rotating, Supplemental Fig. 1C). To distinguish between planar and rotating waves, we used the circular-circular correlation coefficient across three points of the array (shown by dark circles in Fig. 3B). Each of these coefficients had a distinct sensitivity profile to rotational and planar waves. Thus, the combination of the three ρ_c_ values provided a fingerprint-like signature that could be used to accurately classify wave types (planar vs rotating) and identify their wavelength and direction (Methods). Simulations were used to quantify ρ_c_ around the three points for different types of planar and rotating waves (see Methods, Supplemental Fig. 1D,E). The set of coefficients computed for the LFP data was then compared with the simulation sets to classify waves based on which set was closest number of planar vs rotating waves observed across all four arrays. Overall, rotational waves were more common than planar waves for all frequency bands in all task epochs. An example theta rotating wave (Subject 1, right dlPFC, memory delay) is shown in Movie 2 (Supplementary Materials). This method was not constrained by the exact choice of points on the array (Methods). Analyzing ρ_c_ values around other points on the array (Supplemental Fig. 2A) showed similar results.

Our arrays were relatively small (3×3mm), meaning that it was possible, if not likely, that our arrays were capturing pieces of a larger rotating wave structure. A rotating wave will show different spatial wavelengths (i.e. the spatial distance from one wave peak to the next) depending on the position of the center of the rotating wave relative to the array. As one moves away from the core of the rotating wave, the spatial wavelength increases. We quantified the incidences of waves of different spatial wavelengths (Fig. 3D). For all frequency bands, we observed a greater incidence of waves with longer wavelengths. This suggests that, indeed, we were capturing pieces of a larger rotating structure. We will return to this point later.

### Waves travelled in preferred directions

The waves did not travel in random directions. To check if there were preferred directions of travel along each array, we leveraged a property of the circular-circular coefficient. The coefficient could discriminate between waves in opposing directions (red and blue regions respectively, Fig. 3A) along a particular axis. Waves directed towards the positive half (red arrows, Fig. 3A) had ρ_c_ >0 and vice versa (blue arrows). The orientation of the axes splitting the positive and negative regions depended on the net direction of the rotation map chosen and hence differed with the chosen point on the array (Fig. 3A, left vs right, the chosen point shown in yellow). We chose points such that we could split the array into polar segments of around 10-15 degrees each. We confirmed these properties with simulated data (see Methods). To test for statistical significance of the correlation obtained, we calculated a ρ_c_ threshold beyond which the phase organization exceeded chance (p<0.01, ρ_c_ >0.3, random permutation test). This approach allowed us to measure wave directions but did not discriminate between planar and rotating waves.

Fig. 3E shows the placement of one of the arrays relative to the principal and arcuate sulci, with polar histograms of observed wave directions overlaid. Some wave directions were more common than others. This was consistent across the frequency bands. We reoriented the data from all four arrays such that their most preferred direction (for each frequency band) was along the horizontal axis (Fig. 3F). The wave counts in each direction were then averaged across trials. Clear directional preferences were seen across all arrays and all frequency bands (theta to beta). Note that the directional preference remained consistent across trials epochs including the baseline (dashed lines). In other words, there were preferred “default” directions. Task performance increased or decreased the probability of waves traveling in those directions. A degree of bimodality was also observed for all frequencies. In addition to the preferred direction at 0° (by definition), there was a secondary preference for directions around 180°. Thus, there was typically a preferred *axis* of wave propagation (dashed arrow, Fig. 3F) with an additional preference for one direction over the other along that axis.

### Task-related changes in wave direction

Though task demands did not change the preferred axis of wave motion, we found that it could alter the balance between opposite directions along that axis. To determine this, we again used the circular-circular correlation analysis. For this purpose, we examined correlation values (ρ_c_) with the rotation map around the point that showed the maximum number of waves for each array, adjusting such that the task-enhanced direction had positive ρ_c_. This approach included both planar and rotating waves. Oppositely directed waves showed opposite signs.

Figure 4A shows the distribution of ρ_c_ values, averaged across all trials and all four arrays in both animals. Results are shown separately for three frequency bands, and for three task epochs (green lines), with each compared to the pre-trial baseline (red lines). The ρ_c_ at each time instant was classified as a wave if it exceeded the value expected by chance (indicated by shaded areas of Fig 4A, see Methods). Alpha and theta waves showed unimodal ρ_c_ histograms centered around zero during baseline and delay epochs. Presentation of the sample or test-array shifted the histogram peak out of the shaded area (indicating increase in wave incidence) but did so in one direction over the opposite. This could be seen in theta and alpha frequency bands (Fig 4A).

**Figure 4:**
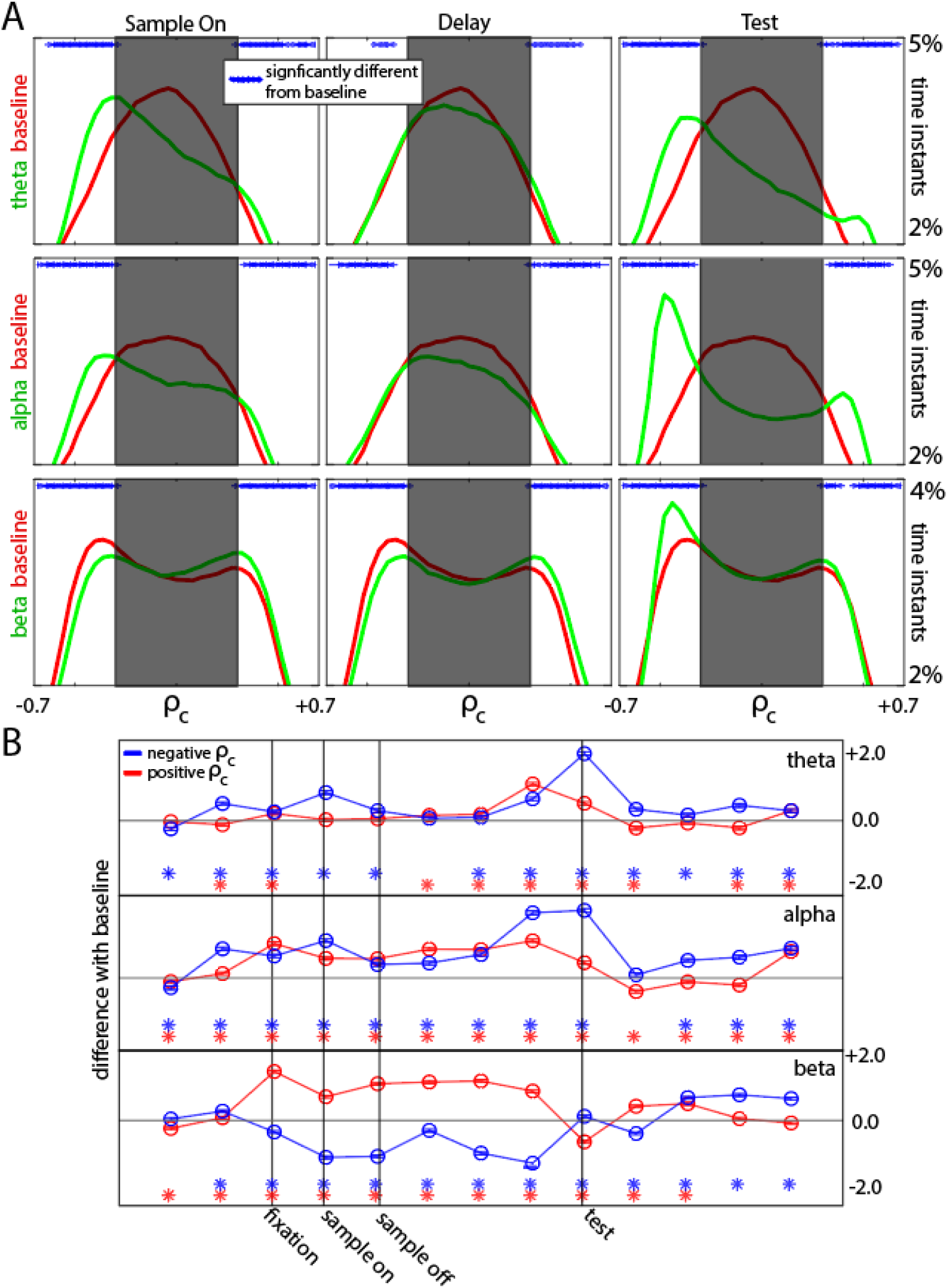
(A) Histograms quantifying the correlation values seen on average in each trial during the sample, delay, and test-onset intervals (all 0.5 s in length), combined across arrays. A correlation value to the right of the shaded region (positive) denotes waves in a particular direction, while the left means the opposite direction. The shaded region denotes the “chance zone” where no conclusion regarding wave direction can be made. For each frequency range (each row), the red histogram corresponds to the correlations observed in baseline conditions (0.5 s pre-fixation), while the green histogram corresponds to the correlations observed in that epoch (0.5 s). The blue dots denote if the two are significantly different from each other (p<0.01). (B) Quantification of the difference between the green and red curves in (A) for all 0.5 s intervals during each trial. The red line shows difference from baseline for the positive wave direction, while the blue line for the negative wave direction.

Beta waves showed prominent bidirectionality throughout the trial (Fig. 4A, bottom row). During both the baseline (red line) and task performance (green line), the distribution of ρ_c_ values for waves in the beta band were bimodal with “bumps” on the ends of the distributions (outside the “chance zone”). This indicates waves in opposite directions. Relative to baseline, during task performance the “bump” on one end of the distribution rose while the other lowered, indicating an increase in waves in one direction and a decrease in the opposite. Baseline ρ_c_ values (red lines) for the beta band skewed toward the negative direction indicating a baseline default bias for waves to travel in that direction. During the sample and delay epochs (green lines), waves skewed more toward positive ρ_c_ values). After test screen presentation and the monkey’s behavioral response, this reversed back to a skew toward negative ρ_c_ values, the baseline direction opposite of that seen in the task.

This is illustrated in more detail in Figure 4B. It shows changes in wave direction bias (relative to baseline) over time. For theta and alpha waves, negative ρ_c_ values were significantly increased from baseline during the sample (blue line, Fig 4B) and especially after the test screen appeared. Positive ρ_c_ values also increased intermittently for these two frequency bands. The beta band showed the most consistent changes in wave direction preference with task performance (Fig. 4B, bottom row). During sample presentation and the memory delay, there was an increase of positive ρ_c_ values and a decrease in negative ρ_c_ values indicating a consistent shift toward waves flowing in one direction over the other. After test screen presentation and the animal’s behavioral response (i.e., post-test), this shift ended. There was a decrease in positive, and an increase in negative ρ_c_ values resulting in the mix of the two directions seen during baseline.

For this representation, we adjusted the coefficient points such that the enhanced wave direction had positive ρ_c_ for all frequencies. However, it is important to note that although the waves in different frequency bands had similar preferred direction *axes* (Fig. 3F), this did not mean that the waves in different bands traveled in the same directions at the same time, as part of a multiband wave. Supplemental Fig. 3 shows the correlation coefficient calculated around the central point (4,4) for the left dlPFC array of Animal 2. As can be seen in the histograms (Supplemental Fig. 3A), theta waves preferred the negative direction during baseline, sample presentation and delay (higher histogram bump towards negative ρ_c_ values). By contrast, beta waves preferred the opposite direction, i.e., the higher histogram bump was towards positive ρ_c_ values. This is also evident in the quantification of the directional increase during task performance from baseline (Supplemental Fig. positive values were enhanced for beta waves.

### Inferring the larger wave structure from observed dynamics

For the analyses above, we aligned the waves along the preferred axis/direction of the waves for each array. In reality, the waves flowed in different anatomical directions across each individual array. This was likely due to different placement of each array relative to a larger wave structure. We used features of rotating waves to infer that larger structure.

Rotating waves create a heterogeneous vector field with the following features (illustrated in Supplemental Fig. 1D,E): 1. The local movement of waves from the same rotating structure will flow in different directions in different areas around the center of rotation. 2. Rotating wave organization decays as one moves away from the center causing the wave to lose structure. 3. The spatial wavelength of the wave (the spatial distance from peak to peak) grows as one moves away from the center of rotation. Thus, shorter wavelengths on the array indicated that the rotating wave center was closer to the center of the array and vice versa. We also used our correlations coefficients to quantify the exact direction and type of wave pattern (Supplemental Fig. 2B,C). We estimated the locations of the rotating waves using these two metrics. The three frequency bands had similar wave directions and curvatures. However, the beta band tended to show the highest traveling wave counts (e.g., in Fig 4A). Thus, to make better inferences, we focused on the beta band.

The results were consistent with our arrays capturing parts of two oppositely rotating waves. The rotating waves seemed to be located similarly in each hemisphere for both subjects relative to anatomical brain landmarks (the arcuate and the principal sulci). The locations of the Fig. 5A shows the spatial wavelengths recorded for Animal 1, along with the directional histograms for beta-band waves. The array in the right hemisphere recorded more short-wavelength waves than the array in the left hemisphere. The result is a wider ρ_c_ histogram spread and stronger directional differences than the left hemisphere array. This would suggest that the rotating wave centers were closer to the array in the right hemisphere, i.e. the recording array captured more of the rotating waves in this hemisphere. Based on the wave features described above (shown as bar graphs), Fig. 5B shows hypothesized locations of the larger rotating wave for each rotational direction relative to anatomical landmarks in each hemisphere (see Methods).

**Figure 5:**
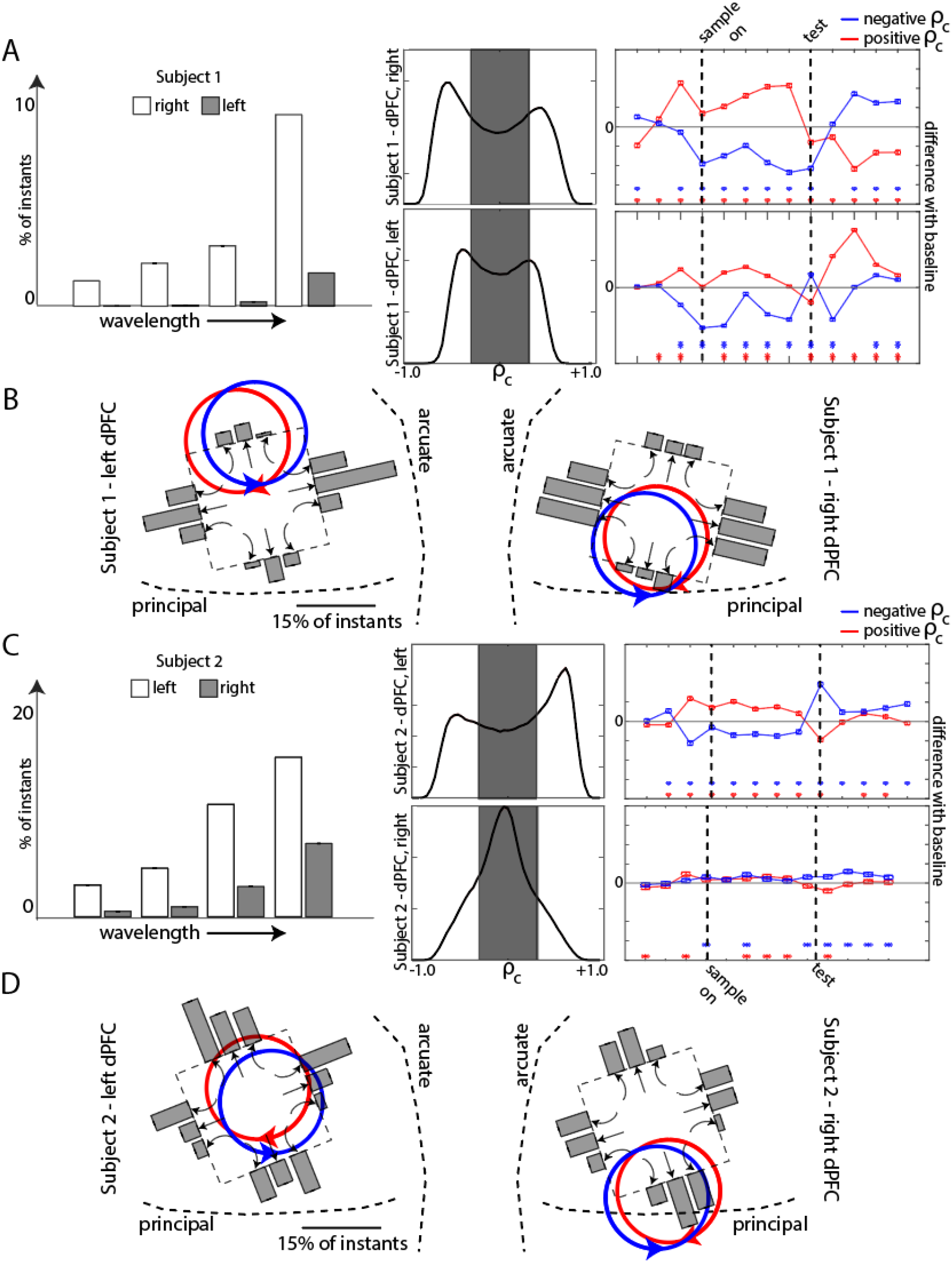
(A) Quantification of the differences observed in beta-band waves between the left and right hemispheres of Subject 1 with regard to spatial wavelengths observed (left), correlation coefficients (center), and wave directional differences (right). (B) Quantification of the levels of different wave types (bar graphs, with the wave type indicated through arrows below the corresponding bar graph) on each array – overlaid with respect to the array position in the brain – for both left and right hemispheres of Subject 1. The red and blue rotating directions indicate the inferred locations of the opposite rotating waves based on the pattern of different wave types observed. (C,D) Same as (A) and (B) but for Subject 2.

Fig. 5C shows similar analysis for Animal 2. In this case, the array in the left hemisphere showed significantly stronger effects than the right, indicating the position of the left hemisphere array was closer to the center of rotation. For example, the array in the left hemisphere of Animal 2 recorded the highest number of short spatial wavelengths. Thus, it can be hypothesized that the rotating wave covered the most array area compared to the other three arrays.

There was a correlation between spike rate and rotating wave organization. Spike rates were higher closer to the center of the rotating wave (Supplemental Fig. 4). Shorter wavelengths indicate that the center of the rotating wave is closer to the array (Supplemental Fig. 4A). We found that shorter wavelengths were associated with higher spike rates (Supplemental Fig. 4B). This indicates that rotating waves modulate spike rates.

## Discussion

We found that 4-30 Hz (theta, alpha, and beta band) oscillations in the lateral prefrontal cortex organize into traveling waves. Rotating waves outnumbered planar waves. Average spike rates were higher near the rotating wave centers. Before the trial began, traveling waves had a preferred orientation but tended to be bidirectional, flowing in opposite directions randomly. After behavioral trial initiation, there was an increase in waves in one direction. This was especially evident during presentation of a to-be-remembered sample and subsequent presentation of a test screen of two stimuli. Notably, in the beta band, there was a persistent increase in the flow of waves in one direction while waves in the opposite direction decreased during the delay. Our animals were well trained on the task showing correct responses in 87% of the trials. This did not give us enough statistical power to compare traveling wave characteristics in correct vs error trials. Future studies will be needed to address this.

### The neural infrastructure of traveling waves

The existence of traveling waves provides insight into the underlying circuitry. They can be explained by short-range lateral interactions between weakly-coupled oscillators (12). This creates phase gradients, i.e. time lags, between successive regions resulting in a sequential movement of a peak of activity.

We found that the waves flowed back and forth in opposite directions. This could be explained by activation of different subsets of neurons with different wave direction preferences. Increasing activation of one network could inhibit the other thus biasing flow in one direction (24), as we observed during task performance especially in the beta band. Removal of activation from this network and the corresponding release of inhibition from the other network could explain our observation of a decrease in preferred, and an increase in the non-preferred, wave direction after the trial ended.

The mechanism of generating rotating waves is a subject of extensive theoretical research (25). A rotating wave is different from a planar or radial wave. Rotating waves create a heterogeneous vector field, i.e., a different phase-gradient depending on where one is recording (26). It has altered curvature along the arm of the wave. That is, different points along the wave have different curvature, hence different speeds (27), rendering it an extra degree of freedom when compared to planar waves. Rotating waves are formed when spatial heterogeneities lead to the formation of phase-singularities. This heterogeneity is mostly caused by a core circuit with altered excitation/inhibition levels around which the wave starts to rotate (28). How this altered core forms varies from field to field (29,30), and in the context of neural mechanisms, remains an open question.

While most studies have reported planar travel waves; a few, like ours, also found rotating waves (26,31). Rotating waves span wide expanses of human cortex during sleep spindles, appear in visual cortex of turtles following sensory stimulation, or spontaneously in rodent cortical slices (visual cortex). But as we demonstrated in our analysis, the location of the recording arrays relative to the wave affects observations. Rotating waves also tend to drift around in space, further complicating the consistent capturing of the rotating center on the recording array (28). This is consistent with analyses suggesting that some planar waves might be incomplete observations of parts of rotating waves (32). More accurate assessment of the relative proportions of rotating vs planar waves can be made with larger recording arrays.

### Functional role of rotating/traveling waves in working memory

One advantage of traveling waves over synchronous oscillations (standing waves) is in maintaining current network status/information. Standing waves result in time periods when all neurons in a network or subnetwork are turned “off”. By contrast, traveling waves ensure that a subset of a given network is always “on” (32). Maintaining “on” states is particularly pertinent for holding items in working memory. Indeed, a recent modeling study shows that segregating memories and maintaining them in neural modules is more robust and stable when the oscillations are phase-shifted among neural ensembles, i.e. manifest as traveling waves (33). Although traveling waves were dominant (Supplemental Fig. 5A), “standing-wave”-type oscillations were also observed, albeit in much lower numbers. It is important to note that a perfect standing wave (no phase variance) is unlikely. Oscillations could be considered “standing” when there was no clear phase gradient and the phase variance was lower than for a traveling wave (Supplemental Fig. 5).

Traveling waves may also play a role in a fundamental cortical function: predictive coding. The brain continually generates predictions of immediately forthcoming inputs, to prevent processing of predicted (thus uninformative) inputs, preventing sensory overload (34). Unpredicted/new inputs are allowed to pass as “prediction errors”. A model of the cortex built by VanRullen and colleagues (22) shows how alpha-band traveling waves carry “priors” from higher to lower cortex in the absence of sensory inputs. Sensory inputs evoke traveling waves that flow in the opposite direction.

Stimulus-evoked traveling waves can add information about time and recent activation history to the network. Muller et al. 2018 (19) suggest that traveling waves allow temporal reversibility: One can decode the history of a pattern from the observed spatiotemporal structure. A given pattern of network activations sets up a unique pattern of traveling waves. This temporal sequence can be decoded to track the elapsed time and recent activation patterns. Elapsed time was not behaviorally relevant in our working memory task; animals were cued when to respond. But relevant or not, the brain keeps track of time. A common example is PFC spiking ramping up near the end of a fixed delay interval in expectation of the forthcoming decision or behavioral response (9,35–37). Recent work has shown PFC neurons track time via neurons that spike at particular times during a memory delay (38). Traveling waves may provide a mechanism of such tracking, through phase-based temporal encoding – i.e. assigning a unique phase map to each instant of time. Tracking time potentially allows neurons to be prepped for future events rendering the cortex with predictive abilities. We did not observe any changes in traveling wave with different sample items. This is consistent with theories that traveling waves have “meta” network functions independent of the items or other content being processed as mentioned above.

We found higher firing rates closer to the center of the rotating wave. This modulation may provide computational advantages over its planar counterparts. Rotating waves have been proposed to play a role in organizing and inducing neural plasticity. The repetitive dynamic of the wave caused by the rotating arm coming around in precise time-intervals can create a structure of timing differences in neural activation that induce spiking timing-dependent plasticity (STDP) in specific subsets of a network. A human ECOG study has shown rotating waves during sleep-related memory consolidation with timing differences in STDP range (26). They showed that the coordinated organization of traveling waves is vital in maintaining plasticity. Short-lived (< 1 sec) increases in synaptic weights induced by spiking may aid in WM maintenance (39). Thus, the waves we observed in the PFC may play a role in not only inducing this plasticity but also periodically refresh them (with spiking) so that memories can be held beyond the time constant of the short-term plasticity (40).

### Summary

Neural oscillations in the low frequency ranges have been implicated in a variety of functions, including working memory, attention, and predictive coding (41-46). Here, we show that they manifest in the prefrontal cortex as traveling waves that change with task performance. Traveling waves have multiple characteristics – such as direction, phase organization, speed - all of which can serve specific functions, making these waves a potentially powerful computational tool. Given their functional advantages, a greater understanding of traveling waves should lead to a greater understanding of cortical function.

## MATERIALS AND METHODS

### Subjects, task and LFP recordings

The nonhuman primate subjects in our experiments were two adult males (ages 17 and 8, for Subject 1 and 2 respectively) rhesus macaques (*Macaca mulatta*). All procedures followed the guidelines of the Massachusetts Institute of Technology Committee on Animal Care and the National Institutes of Health (protocol 0619-035-22, approved by MIT’s Committee on Animal Care on 6/23/21).

Subjects performed a delayed match-to-sample working memory (WM) task. They began task trials by holding gaze for 500 ms on a fixation point randomly displayed at the center of a computer screen. A sample object (one of eight randomly chosen) was then shown for 500 ms at the center of the screen. After a 2 s delay, a test object was displayed. The monkeys were required to saccade to it if it matched the remembered sample. Response to the match was rewarded with juice. All stimuli were displayed on an LCD monitor. An infrared-based eye-tracking system (Eyelink 1000 Plus, SR-Research, Ontario, CA) continuously monitored eye position at 1 kHz.

The subjects were chronically implanted in the lateral prefrontal cortex (PFC) with two 8×8 iridium-oxide “Utah” microelectrode arrays (1.0 mm length, 400 μm spacing; Blackrock Microsystems, Salt Lake City, UT). Signals were recorded on a Blackrock Cerebus. Arrays were implanted bilaterally, one array in each dorsolateral PFC. Electrodes in each hemisphere were grounded and referenced to a separate subdural reference wire.

LFPs were band-passed from 0.5-300Hz and sampled at 1kHz. There were two filter stages, first a real-time analog filter in the amplifier, then a symmetric digital filter offline in software:

a. Analog filter: High-pass: 0.3 Hz 1^st^ order Butterworth Low-pass: 7.5 kHz 3^rd^ order Butterworth
b. Digital filter: Low-pass: 300 Hz 3^rd^ order Butterworth

All correctly performed trials were included in analyses. All preprocessing and analysis were performed in Python or MATLAB (The Mathworks, Inc, Natick, MA). For the power analysis, the resulting signals were convolved with a set of complex Morlet wavelets.

### LFP spatial phase maps

The raw LFP traces were filtered in the desired frequency range, using a 4^th^ order Butterworth filter, forward-reverse in time to prevent phase distortion (see MATLAB function *filtfilt*) and interpolated for missing electrodes. Out of 28 sessions, five had missing electrodes (one electrode missing in 3 sessions, three electrodes missing in 2 sessions). Linear interpolation was done to account for the missing data. Overall as these numbers were a minority, our statistics were not affected. A Hilbert transform was used to obtain the analytical signal for each electrode. The phase of each electrode for the 8×8 array is called the “phase map” for that time instant. These phase maps (unsmoothed) were checked for gradients to identify traveling waves.

### Traveling wave identification through linear distance maps

Traveling waves were identified at an instant in time when spatial correlation between the phase map at that instant and a Euclidean distance map template (Supplemental Fig. 1A) in a particular direction exceeds a certain threshold. Pearson correlation coefficients were computed over the full 2-dimensional array of values. The threshold was decided through a shuffling permutation procedure (26).

The distance map was made for each quadrant of the array, in the particular direction being evaluated (Supplemental Fig. 1A shows the phase map for a diagonal wave for one quadrant). The distance maps were made such that they always increased towards the edges. So, a wave entering the array from an edge would encounter a direction map in the opposite direction – and hence show a negative correlation, while a wave leaving the array would produce a positive correlation, as it increased towards the edges. A wave thus was counted when one quadrant showed a positive correlation with the phase map, while another showed a negative correlation. Then a wave was counted to have passed from the negative quadrant to the positive quadrant. This method thus identified a wave component in a particular direction – from one quadrant to the other. Four quadrants yielded a total of 12 directions. This quadrant-based method allowed identification of wave fragments even when waves did not have uniform spatio-temporal structure throughout the whole array. So, when one wave passed, multiple direction components could be recorded. This method was verified using simulated waves.

Only positive phase values were considered in this method, so as to identify consecutive bursts of traveling wave instances. That is, when two consecutive waves passed, this method showed two peaks (two positive cycles), instead of one continuous high value for both waves. As a consequence, however, this method was restricted to identifying waves that had a long spatial wavelength when compared to the size of the array, to allow for instances where only positive phase values are on the array at one time. This approach was only used for planar wave detection. This approach is consistent with earlier studies that suggest that the spatial wavelength of traveling waves is long compared to the LFP recording array size (23).

### Wave speed calculation

Wave speed at a time instant was calculated from the phases (*p*) by dividing the temporal frequency (*∂p/∂t*) at that time with the spatial frequency (*∂p/∂x*) (16). The gradients obtained were averaged across electrodes to get the net wave speed for that time instant. Essentially, this calculated, how quickly the oscillation cycle progressed in space versus in time. For a fast wave, points away from each other on the array will reach the same phase within a shorter time period, i.e. a shallow spatial gradient (same *∂p* for a larger *∂x* in a certain *∂t*), when compared to a slower wave. Wave speed was calculated only for waves of long wavelength (majority of wave instants) as spatial frequencies for short wavelength waves could vary along the array.

### Shuffling procedure

To ensure that the probability of detecting traveling waves exceeded that expected by chance – we performed a random shuffling procedure to establish a threshold for the correlation coefficient – beyond which a traveling wave was counted. This was done by shuffling the phase values on the array randomly (with 25 different types of random permutations) and calculating the correlation coefficient. The 99^th^ percentile of the resulting distribution of coefficient values determined a threshold (0.3) above which the correlation exceeded chance.

It is important to mention here that characterizing traveling waves through quadrant-based methods or patch-based methods (47) for small arrays may lead to erroneous conclusions especially for waves with variable wavelengths or with variable local dynamics. For that purpose, we used our quadrant-based approach only for long wavelength waves which had a consistent structure on the array. To classify rotating wave patterns, we shifted to circular statistics using the whole 8×8 array.

### Circular-circular correlation coefficient – planar waves, rotating waves, and net wave directionality

We used circular statistics to identify wave patterns. Circular-circular correlation coefficients have been used to classify waves in earlier studies (26). In this study, we demonstrate that the use of circular-circular correlation coefficients can be extended to distinguish between a spectrum of wave types.

The circular-circular correlation coefficient reports the spatial gradient similarity between two phase maps that are adjusted to account for circular phase values. Adjusting for circular values reorganizes the circular phases by subtracting the circular mean (so the mean becomes 0, and phases range from -π to π), and then taking the sine, such that increasing values now directly translate to increasing oscillation phase. In our case, we checked spatial gradient similarity between a phase map obtained from the LFP recordings with a rotation map template. The rotation map expressed the polar angle of each point on the array relative to a particular origin point (Supplemental Fig. 1B). For the signal phase (*φ* at each array coordinate [a,b]) and the rotation angle (θ) around the chosen point, the circular-circular correlation coefficient thus was (26):

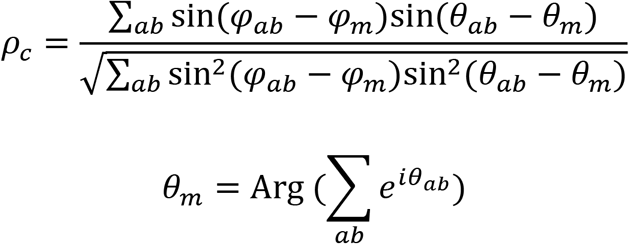

#### Net direction and bisecting axes

A rotation map around a particular point has a particular *net direction* that can be revealed by plotting the phase maps, adjusting for circular values (i.e. subtracting the circular mean and sine transforming). The overall gradient of a phase map can best be appreciated by looking at the circular-adjusted maps. Supplemental Fig. 1B shows the net directions (solid lines) for rotation maps around three points. This was obtained by summing the gradients in phase across the array as complex vectors – the resultant vector being the net direction. It can be appreciated from this figure that while actual phase values may range anywhere between -pi to pi, the circular adjustment reorients all values from – 1 to 1. A positive coefficient value could be obtained by multiple types of waves – rotating waves centered near the chosen point in the direction of the rotation map, or planar waves moving towards the map’s *net direction*. Of course, opposite directions for both would yield negative values. The orthogonal direction would result in a coefficient close to zero – which creates the *bisecting axis*. This approach thus allowed us to classify wave directions into positive and negative halves separated by this bisecting axis (Fig. 3A) – taking into account both rotating and planar waves. The axis was dependent on the rotation map, which varied with the chosen point.

Similar to the earlier wave classification – a correlation threshold was chosen below which no conclusions regarding wave directions could be made. This threshold was identified through shuffling the electrodes and ensuring phase gradients did not appear by chance. The 99^th^ percentile of the resulting distribution of coefficient values determined a threshold (0.3) above which the correlation exceeded chance. This method allowed us to categorize waves into two opposite directions (Fig. 4) and analyze how the directions changed during task performance. For Fig. 4, rotation maps were chosen for each array in a way that maximized the number of waves observed for that array, to provide the greatest power to discriminate between opposing directions of wave motion (Subject 1: right array – (4,4), left array – (8,4); Subject 2: right array – (4,1), left array – (4,4)).

#### Planar vs rotating waves

To distinguish between planar and rotating waves, however, just one rotation map would not suffice, as the values for a rotating wave and a planar wave with the same net direction could be similar. The coefficient value using a rotation map around a particular point essentially quantifies the gradient vectors seen in that phase map. Considering, for example, a diagonal planar wave from the bottom left to the top right (Supplemental Fig. 1C, left). The central point (4,4) has a net direction along that diagonal and hence records a high value around 0.9. The point (4,1) has a net direction along the horizontal and records an intermediate value around 0.64. Now, let’s consider a wave that rotates from bottom left to top-right (Supplemental Fig.1C, right). The circular-reoriented phase maps for the two waves are shown below. It is evident that the increase toward the top right is still maintained causing the coefficient around (4,4) to remain the same. However, the increase toward the horizontal (towards the right) is less when compared to the diagonal wave case. This reduces the coefficient around (4,1) to 0.52. Thus, while one coefficient could not distinguish the types of waves, a combination of two coefficients in this case was able to do so.

#### Wave classification, wavelength and wave structure

In this way, we show using simulations that three coefficients, around points (4,4) – diagonal axis, (1,4) – horizontal axis, and (4,1) – vertical axis, could be leveraged together to distinguish planar and rotating waves. We created simulated datasets for these three coefficients. Waves were simulated using the following equations (26):

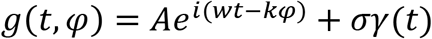

where *φ* was the input phase map (rotational or planar), *w* was the temporal frequency and *k* was the spatial wavenumber (1/wavelength). The second term was a Gaussian white noise term, with zero mean and standard deviation *σ*.

Using the above simulation equation, we created a dataset comprised of 32 types of planar waves in all directions, each separated by around 12 degrees, and 40 types of rotating waves of different curvatures and wavelengths (*k* ranging from 0.1 to 0.9) (Supplemental Fig. 1E). As evidence for the property described in the section above, if one monitors the (4,4) red curve in this figure, it is clear that values change signs between waves directed towards the bisecting axis (top-left to bottom-right diagonal in this case).

A long spatial wavelength (inverse of wavenumber *k*) was chosen for planar waves simulations (Supplemental Fig. 1D, top), as their wavelength remains constant throughout the wave band. But, rotating waves show different wavelengths at different portions of the wave (Supplemental Fig. 1D, bottom). As one moves away from the rotating center (white circle), the wavelength increases. Hence, multiple wavelengths were considered while creating the datasets for rotating waves. It is also evident from Supplemental Fig. 1E that, in the case of rotating waves, as spatial wavelength increases (lower *k*) – all coefficients start to decrease – proving that the wave starts to lose spatio-temporal structure away from the rotating center. An average of 20 simulations (with distinct realization of Gaussian noise) were used to calculate the coefficient for each type of wave. We ensured at least one coefficient value always remained above the chance threshold (0.3) to ensure sufficient wave structure (which is essentially why three points where needed).

We compared the values of three coefficients obtained from each time instant of the working memory trials with the simulated datasets to then automatically classify the type of wave observed, based on the Euclidean distance between a wave type and the observed phase map. Specifically, the three ρ_c_ values for each wave type could be represented as a point in three-dimensional space. The Euclidean distances between the corresponding point for an observed phase map and all wave types were computed. The phase map was classified into the wave type from which it was the closest. Coefficient matching with three different maps, allowed for greater accuracy with lesser chances of misclassification.

#### Choice of array points for coefficient calculation

Our results were not constrained by the exact choice of points on the array around which the circular-circular correlation coefficient was to be calculated (Supplemental Fig. 2A). Our intuition for choosing (1,4), (4,1) and (4,4) were based on trying to capturing the whole spectrum of wave types. (1,4) was able to differentiate waves that traveled roughly towards the top-half of the array vs those that traveled towards the bottom-half (Supplemental Fig. 1B). However, (1,4) had a “chance-zone” along the horizontal axis, i.e. it could not identify waves that traveled in horizontal directions. For that purpose, another coefficient – around (4,1) - was necessary to capture waves that traveled laterally along the array. An extra point (4,4) was added to identify rotating waves that naturally occurred with their center on the array.

#### Inferring rotating wave locations

In Fig. 5B, the locations of the rotating wave directions were inferred based on the levels of the curvature recorded, the histogram spread observed and the number of short-wavelength waves recorded. For example, in Subject1, the right hemisphere array captured a greater incidence of short-wavelength waves – leading to the conclusion that more of the rotation is on the array - when compared to the left hemisphere. The highest wave levels were closer to the ventral side of the PFC, thus suggesting that the part of the rotation that was outside the array was more towards the ventral side. Following this logic, the rotating directions were placed on the array through a subjective estimation based on a combination of all these results.

## Acknowledgments

We thank Jordan G. DeFarias for his technical assistance and Jesus Ballesteros, Andre Bastos, Alex Major, Morteza Moazami, Dimitris Pinotsis, and Jefferson Roy for helpful comments. This work was supported by ONR MURI N00014-16-1-2832, NIMH R37MH087027, and The MIT Picower Institute Innovation Fund.

## Author Contributions

EKM designed the experiments. SLB and ML performed the experiments. SB conceived and performed the analysis. SB, SLB, ML and EKM wrote the manuscript.

## Supplementary Materials

Supplementary Figure 1

Supplementary Figure 2

Supplementary Figure 3

Supplementary Figure 4

Supplementary Figure 5

Movie S1

Movie S2

## References

1. Buzsaki G. Neuronal Oscillations in Cortical Networks. Science. 2004 Jun 25;304(5679):1926–9.

2. Womelsdorf T, Fries P, Mitra PP, Desimone R. Gamma-band synchronization in visual cortex predicts speed of change detection. Nature. 2006 Feb;439(7077):733–6.

3. Fries P. A mechanism for cognitive dynamics: neuronal communication through neuronal coherence. Trends Cogn Sci. 2005 Oct 1;9(10):474–80.

4. Fries P, Fernández G, Jensen O. When neurons form memories. Trends Neurosci. 2003 Mar 1;26(3):123–4.

5. Womelsdorf T, Schoffelen J-M, Oostenveld R, Singer W, Desimone R, Engel AK, et al. Modulation of Neuronal Interactions Through Neuronal Synchronization. Science. 2007 Jun 15;316(5831):1609–12.

6. Bastos AM, Loonis R, Kornblith S, Lundqvist M, Miller EK. Laminar recordings in frontal cortex suggest distinct layers for maintenance and control of working memory. Proc Natl Acad Sci. 2018 Jan 30;115(5):1117–22.

7. Helfrich RF, Breska A, Knight RT. Neural entrainment and network resonance in support of top-down guided attention. Curr Opin Psychol. 2019 Oct 1;29:82–9.

8. Helfrich RF, Huang M, Wilson G, Knight RT. Prefrontal cortex modulates posterior alpha oscillations during top-down guided visual perception. Proc Natl Acad Sci. 2017 Aug 29;114(35):9457–62.

9. Miller EK, Lundqvist M, Bastos AM. Working Memory 2.0. Neuron. 2018 Oct 24;100(2):463–75.

10. Fries P. Rhythms for Cognition: Communication through Coherence. Neuron. 2015 Oct 7;88(1):220–35.

11. Gray CM, Singer W. Stimulus-specific neuronal oscillations in orientation columns of cat visual cortex. Proc Natl Acad Sci. 1989 Mar 1;86(5):1698–702.

12. Kopell N, Ermentrout GB. Symmetry and phaselocking in chains of weakly coupled oscillators. Commun Pure Appl Math. 1986;39(5):623–60.

13. N. Kopell, Ermentrout GB. Coupled oscillators and the design of central pattern generators. Math Biosci. 1988 Jul 1;90(1):87–109.

14. Muller L, Reynaud A, Chavane F, Destexhe A. The stimulus-evoked population response in visual cortex of awake monkey is a propagating wave. Nat Commun. 2014 Apr 28;5(1):3675.

15. Takahashi K, Saleh M, Penn RD, Hatsopoulos N. Propagating Waves in Human Motor Cortex. Front Hum Neurosci. 2011;5:40.

16. Zhang H, Jacobs J. Traveling Theta Waves in the Human Hippocampus. J Neurosci. 2015 Sep 9;35(36):12477–87.

17. Lubenov EV, Siapas AG. Hippocampal theta oscillations are travelling waves. Nature. 2009 May;459(7246):534–9.

18. Sreekumar V, Wittig JH, Chapeton J, Inati SK, Zaghloul KA. Low frequency traveling waves in the human cortex coordinate neural activity across spatial scales. BioArxiv; 2020 Mar.

19. Muller L, Chavane F, Reynolds J, Sejnowski TJ. Cortical travelling waves: mechanisms and computational principles. Nat Rev Neurosci. 2018 May;19(5):255–68.

20. Feller MB, Butts DA, Aaron HL, Rokhsar DS, Shatz CJ. Dynamic Processes Shape Spatiotemporal Properties of Retinal Waves. Neuron. 1997 Aug 1;19(2):293–306.

21. Watt AJ, Cuntz H, Mori M, Nusser Z, Sjöström PJ, Häusser M. Traveling waves in developing cerebellar cortex mediated by asymmetrical Purkinje cell connectivity. Nat Neurosci. 2009 Apr;12(4):463–73.

22. Alamia A, VanRullen R. Alpha oscillations and traveling waves: Signatures of predictive coding? PLOS Biol. 2019 Oct 3;17(10):e3000487.

23. Davis ZW, Muller L, Martinez-Trujillo J, Sejnowski T, Reynolds JH. Spontaneous travelling cortical waves gate perception in behaving primates. Nature. 2020 Nov;587(7834):432–6.

24. Kim R, Sejnowski TJ. Strong inhibitory signaling underlies stable temporal dynamics and working memory in spiking neural networks. 2020;38.

25. Keener JP. A Geometrical Theory for Spiral Waves in Excitable Media. SIAM J Appl Math. 1986 Dec;46(6):1039–56.

26. Muller L, Piantoni G, Koller D, Cash SS, Halgren E, Sejnowski TJ. Rotating waves during human sleep spindles organize global patterns of activity that repeat precisely through the night. eLife. 2016;5:e17267.

27. Bhattacharya S, Iglesias PA. Controlling excitable wave behaviors through the tuning of three parameters. Biol Cybern. 2019 Apr;113(1–2):61–70.

28. Steinbock O, Zykov V, Müller SC. Control of spiral-wave dynamics in active media by periodic modulation of excitability. Nature. 1993 Dec;366(6453):322–4.

29. Panfilov AV, Müller SC, Zykov VS, Keener JP. Elimination of spiral waves in cardiac tissue by multiple electrical shocks. Phys Rev E. 2000 Apr 1;61(4):4644–7.

30. Palsson E, Lee KJ, Goldstein RE, Franke J, Kessin RH, Cox EC. Selection for spiral waves in the social amoebae Dictyostelium. Proc Natl Acad Sci. 1997 Dec 9;94(25):13719–23.

31. Prechtl JC, Cohen LB, Pesaran B, Mitra PP, Kleinfeld D. Visual stimuli induce waves of electrical activity in turtle cortex. Proc Natl Acad Sci. 1997 Jul 8;94(14):7621–6.

32. Ermentrout GB, Kleinfeld D. Traveling Electrical Waves in Cortex: Insights from Phase Dynamics and Speculation on a Computational Role. Neuron. 2001 Jan 1;29(1):33–44.

33. Soroka G, Idiart M. Theta, alpha and gamma traveling waves in a multi-item working memory model. ArXiv210315266 Phys Q-Bio. 2021 Mar 28;

34. Bastos AM, Usrey WM, Adams RA, Mangun GR, Fries P, Friston KJ. Canonical Microcircuits for Predictive Coding. Neuron. 2012 Nov 21;76(4):695–711.

35. Genovesio A, Tsujimoto S, Wise SP. Neuronal Activity Related to Elapsed Time in Prefrontal Cortex. J Neurophysiol. 2006 May 1;95(5):3281–5.

36. Narayanan NS. Ramping activity is a cortical mechanism of temporal control of action. Curr Opin Behav Sci. 2016 Apr 1;8:226–30.

37. Donnelly NA, Paulsen O, Robbins TW, Dalley JW. Ramping single unit activity in the medial prefrontal cortex and ventral striatum reflects the onset of waiting but not imminent impulsive actions. Eur J Neurosci. 2015;41(12):1524–37.

38. Tiganj Z, Cromer JA, Roy JE, Miller EK, Howard MW. Compressed Timeline of Recent Experience in Monkey Lateral Prefrontal Cortex. J Cogn Neurosci. 2018 Jul 1;30(7):935–50.

39. Wasmuht DF, Spaak E, Buschman TJ, Miller EK, Stokes MG. Intrinsic neuronal dynamics predict distinct functional roles during working memory. Nat Commun. 2018 Aug 29;9(1):3499.

40. Mongillo G, Barak O, Tsodyks M. Synaptic theory of working memory. Science. 2008 Mar 14;319(5869):1543–6.

41. Bastos AM, Lundqvist M, Waite AS, Kopell N, Miller EK. Layer and rhythm specificity for predictive routing. Proc Natl Acad Sci. 2020 Dec 8;117(49):31459–69.

42. Buschman TJ, Miller EK. Top-Down Versus Bottom-Up Control of Attention in the Prefrontal and Posterior Parietal Cortices. Science. 2007 Mar 30;315(5820):1860–2.

43. Wutz A, Loonis R, Roy JE, Donoghue JA, Miller EK. Different Levels of Category Abstraction by Different Dynamics in Different Prefrontal Areas. Neuron. 2018 Feb 7;97(3):716–726.e8.

44. Lakatos P, Shah AS, Knuth KH, Ulbert I, Karmos G, Schroeder CE. An Oscillatory Hierarchy Controlling Neuronal Excitability and Stimulus Processing in the Auditory Cortex. J Neurophysiol. 2005 Sep;94(3):1904–11.

45. Womelsdorf T, Fries P. The role of neuronal synchronization in selective attention. Curr Opin Neurobiol. 2007 Apr;17(2):154–60.

46. Voloh B, Valiante TA, Everling S, Womelsdorf T. Theta-gamma coordination between anterior cingulate and prefrontal cortex indexes correct attention shifts. Proc Natl Acad Sci. 2015 Jul 7;112(27):8457–62.

47. Chemali J. Statistical analysis of travelling waves in the monkey primary motor cortex.:17.

