## Supplemental Materials for "Traveling waves in the prefrontal cortex during working memory"

**This PDF file includes:**

Supplementary Figures 1, 2, 3, 4 and 5
Movies S1 and S2

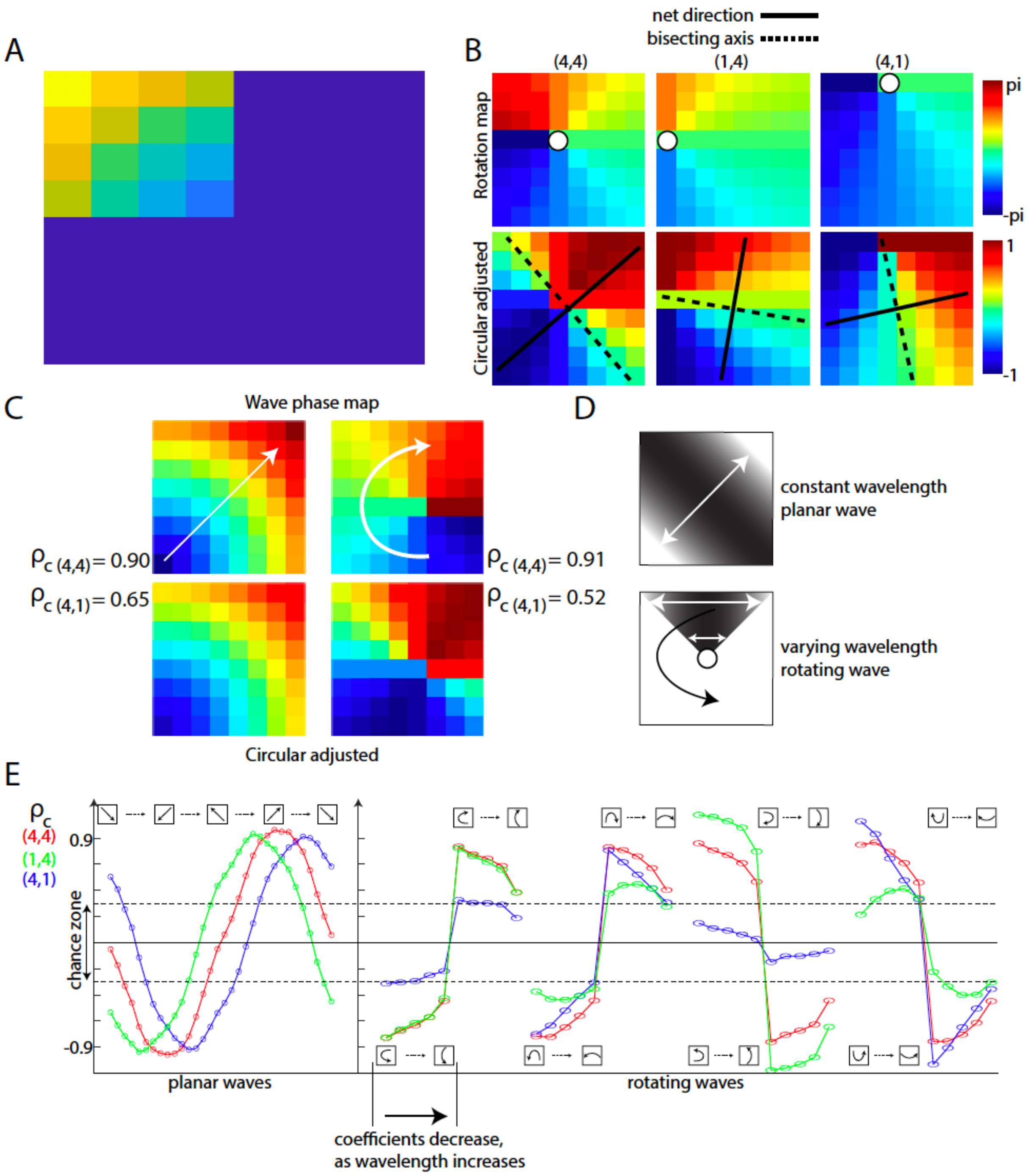


### SUPPLEMENTARY FIGURE 1

Supplemental Fig. 1: (A) Quadrant-based distance map used for traveling wave identification (one quadrant shown for diagonal wave detection). The distance maps would change accordingly for other wave types. (B) Rotation maps around three points on the array (top), adjusted for circular values (mean-centered and sine-transformed; bottom) with the net directions and bisecting axes marked. (C) Example waves (white arrows), with their phase maps (top) adjusted for circular values (bottom) shown with the corresponding coefficient values on the side. (D) Plot illustrating spatial wavelengths for planar and rotating waves. (E) Plot of three correlation coefficient values for different wave types. The region between the dashed horizontal lines denotes the “chance zone”. It was ensured that at least one coefficient for each wave type remained outside this zone.

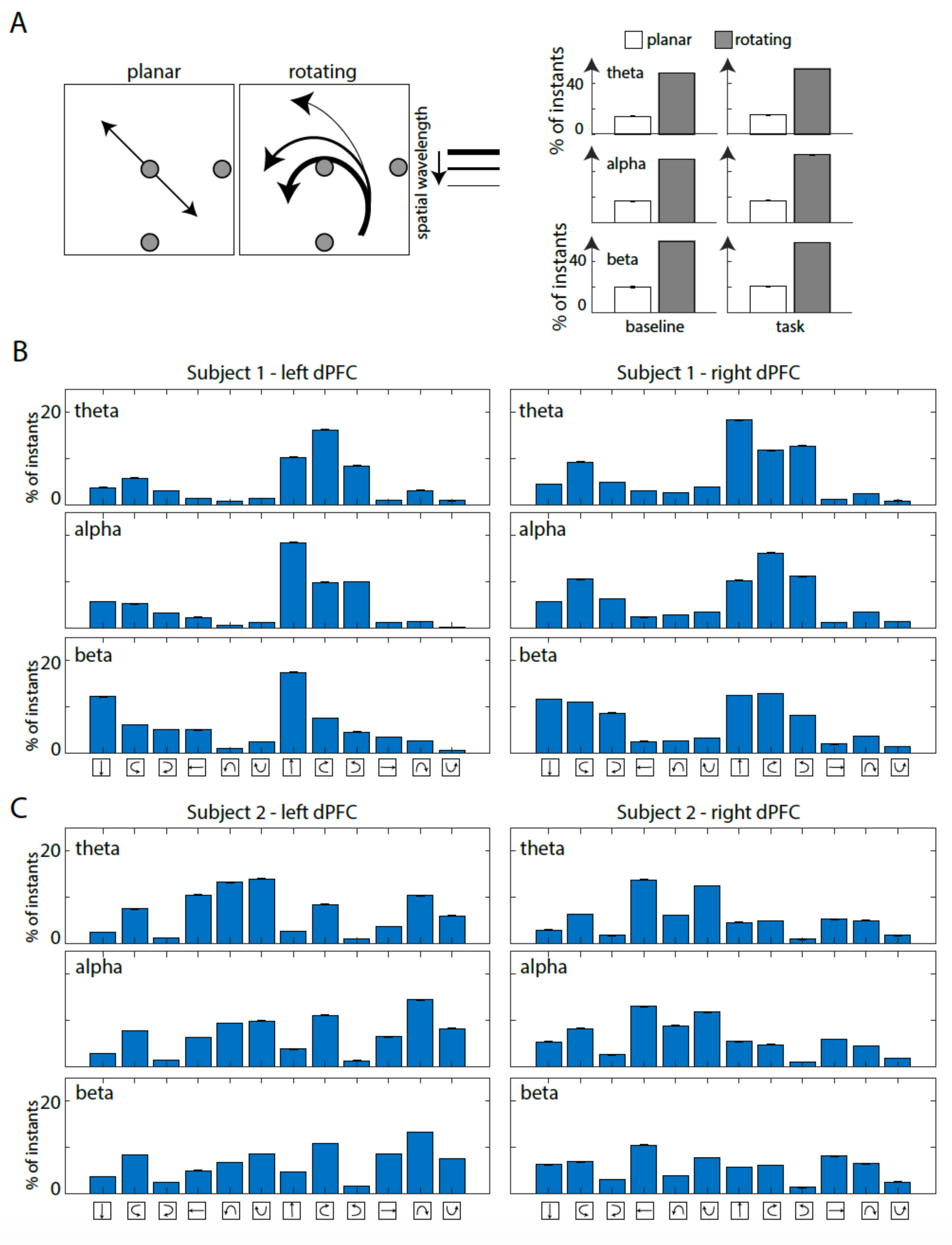


**SUPPLEMENTARY FIGURE 2**

Supplemental Fig. 2: (A) (left) Similar wave classification between planar and rotating waves as Fig. 3, with three different points chosen to distinguish between these wave types. The grey circles denote the points on the array around which the coefficient was calculated. (right) Plots showing the number of time instants in which rotating/planar waves were observed across all arrays and trials in the three frequency ranges. (B, C) Number of wave instants observed for different wave types across frequencies and arrays.

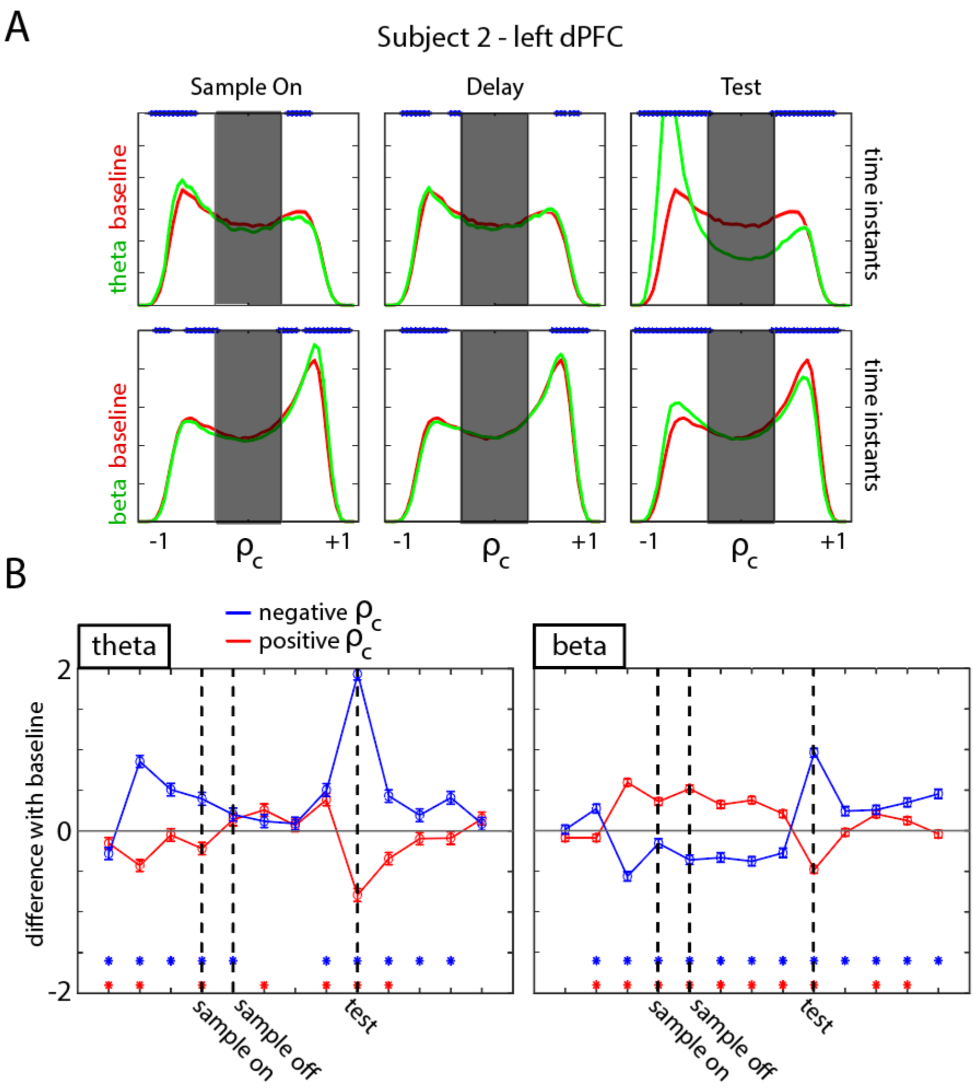


**SUPPLEMENTARY FIGURE 3**

Supplemental Fig. 3: (A) Histograms quantifying the correlation values seen on average in each trial during the sample, delay, and test-onset intervals (all 0.5 s in length), for the left dlPFC array of Animal 2. A correlation value to the right of the shaded region (positive) denotes waves in a particular direction, while the left means the opposite direction. The shaded region denotes the "chance zone" where no conclusion regarding wave direction can be made. For each frequency range (each row), the red histogram corresponds to the correlations observed in baseline conditions (0.5 s pre-fixation), while the green histogram corresponds to the correlations observed in that epoch (0.5 s). The blue dots denote if the two are significantly different from each other ($p<0.01$). (B) Quantification of the difference between the green and red curves in (A) for all 0.5 s intervals during each trial. The red line shows difference from baseline for the positive wave direction, while the blue line for the negative wave direction.

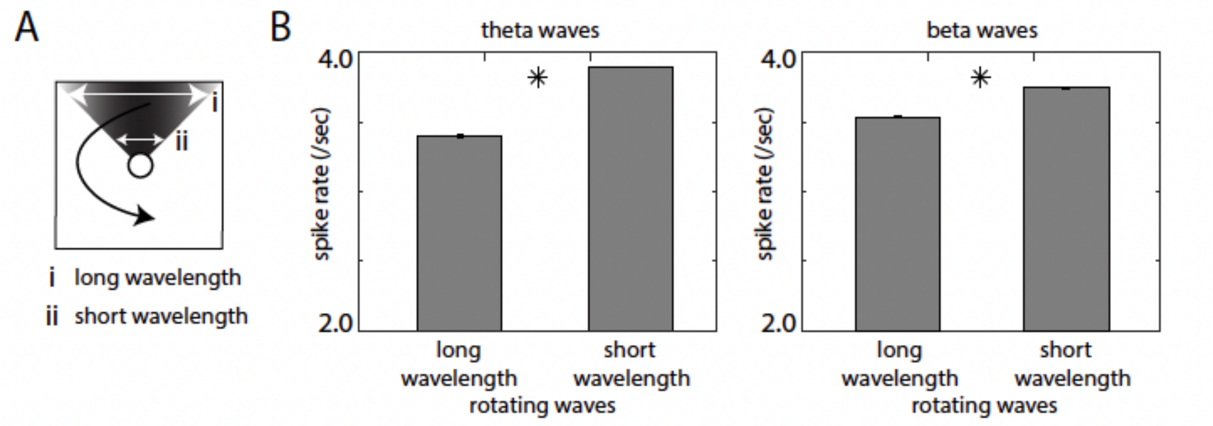


**SUPPLEMENTARY FIGURE 4**

Supplemental Fig. 4: (A) Illustration of a rotating wave with increasing wavelengths at greater distances from the center of rotation (white circle). (B) Spike rates observed for long and short wavelengths, combined across all arrays for theta (left) and beta (right) traveling waves. Star indicates statistically significant difference ($p<0.01$).

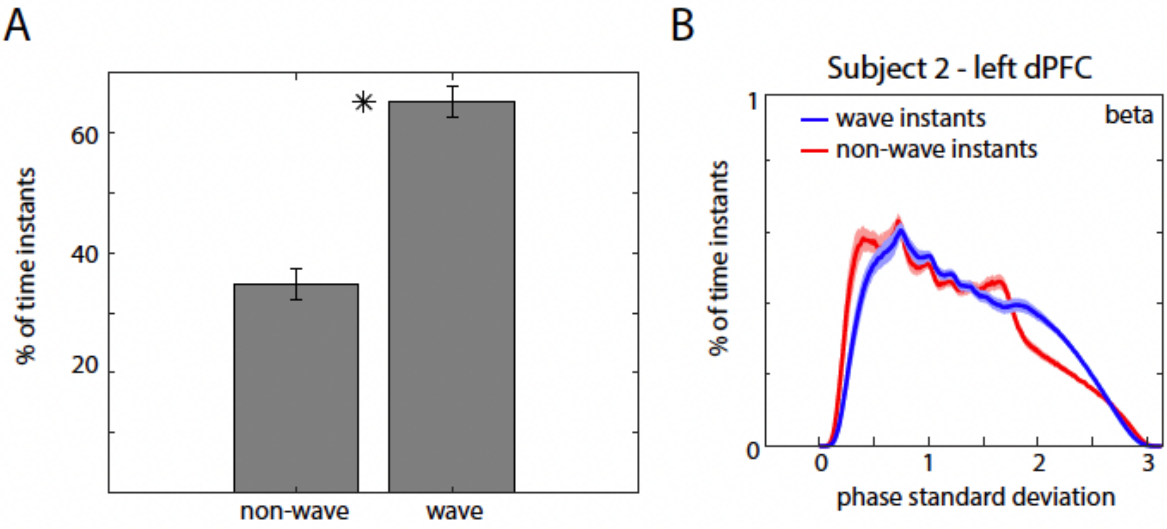


**SUPPLEMENTARY FIGURE 5**

Supplemental Fig. 5: (A) Percentage of wave vs non-wave instants for beta waves combined across arrays. Star indicates statistically significant difference ($p<0.01$). (B) Plot of the phase standard deviation for wave and non-wave instants (normalized probability) observed for beta waves in the left dlPFC array of Subject 2. The red peak to the left of the blue peak corresponds to the standing wave mode, indicating oscillations that were not traveling waves but had a lower phase variance (than traveling waves) on the array.

### SUPPLEMENTARY VIDEOS

**Movie S1**: An example beta-band planar wave on the 8x8 left dlPFC array from a trial in Subject 2, corresponding to the panels in Fig. 1. Each tile marks the LFP oscillation amplitude (warmer colors mean higher amplitude) at that point in time and space.

**Movie S2**: An example of a theta rotating wave on the 8x8 array, during the memory delay period of a trial in the right dlPFC of Subject 1. Each tile marks the LFP oscillation amplitude (warmer colors mean higher amplitude) at that point in time and space.
